# A dual sRNA in *Staphylococcus aureus* induces a metabolic switch responding to glucose consumption

**DOI:** 10.1101/278127

**Authors:** D. Bronesky, E. Desgranges, A. Corvaglia, P. François, C.J. Caballero, L. Prado, A. Toledo-Arana, I. Lasa, K. Moreau, F. Vandenesch, S. Marzi, P. Romby, I. Caldelari

## Abstract

Pathogenic bacteria must rapidly adapt to ever-changing environmental signals or nutrient availability resulting in metabolism remodeling. The carbon catabolite repression represents a global regulatory system, allowing the bacteria to express genes involved in carbon utilization and metabolization of the preferred carbon source. In *Staphylococcus aureus*, regulation of catabolite repressing genes is mediated by the carbon catabolite protein A (CcpA). Here, we have identified a CcpA-dependent small non-coding RNA, RsaI that is inhibited by high glucose concentrations. RsaI represses the translation of mRNAs encoding a major permease of glucose uptake, the FN3K enzyme that protects proteins against damages caused by high glucose concentrations, and IcaR, the transcriptional repressor of exopolysaccharide production. Besides, RsaI regulates the activities of other sRNAs responding to the uptake of glucose-6 phosphate or NO. Finally, RsaI inhibits the expression of several enzymes involved in carbon catabolism pathway, and activates genes involved in energy production, fermentation and NO detoxification when the glucose concentration decreases. This multifunctional RNA provides a signature for a metabolic switch when glucose is scarce and growth is arrested.

## INTRODUCTION

All bacteria require a carbon source, providing energy for their growth, division, and for the synthesis of all macromolecules. Besides, pathogenic bacteria during the infectious process of the host, must cope with hostile conditions such as nutrient deficiency, temperature, oxidative and osmotic shocks, and must overcome innate immune responses. For instance, *Staphylococcus aureus* uses carbohydrates to grow under high NO and anaerobiosis (Vitko et al., 2016). To survive in these complex environments and to counteract the host defense, *S. aureus* produces a plethora of virulence factors. The synthesis of these factors is fine-tuned by intricate interactions between multiple regulators involving both proteins and RNAs (Novick, 2003). Additionally, biosynthetic intermediates, generated from the central metabolism of *S. aureus*, have strong impacts on the synthesis of virulence factors. Besides, several metabolite-sensing regulatory proteins (CcpA, CodY, Rex and RpiR) act as key regulatory factors to coordinate the synthesis of genes involved in metabolic pathways, in stress responses, and in pathogenesis (Somerville and Proctor, 2009; Richardson et al., 2015). Through the adaptation of the metabolism of the bacteria to specific host microenvironment, these proteins contribute to *S. aureus* pathogenesis (Richardson et al., 2015).

Among these proteins, the <u>c</u>arbon <u>c</u>atabolite <u>p</u>rotein <u>A</u> (CcpA) acts as a catabolite regulator, allowing the bacteria to use the preferred carbon source (i.e., glucose) in a hierarchical manner (Seidl et al., 2008a; Seidl et al., 2009). CcpA belongs to the LacI repressor family and binds to a specific DNA sequence, called the *cre* (catabolite responsive element) sequence, which is conserved in many Gram-positive bacteria. Transcription of CcpA is constitutive and the protein is activated through the binding of its co-regulator histidine-containing phosphocarrier protein HPr in the presence of glucose. Inactivation of *ccpA* gene affects the expression of a large number of metabolic encoding genes in a glucose-dependent and -independent manner (Seidl et al., 2008a). Additionally, CcpA has a strong impact on the expression of *S. aureus* virulon. It enhances the yield of the quorum-sensing induced RNAIII, which represses Protein A and various adhesion factors at the post-transcriptional level, and conversely activates the synthesis of many exotoxins. However, CcpA also modulates the transcription of mRNAs encoding Protein A, α-hemolysin (*hla*) and TSST (Seidl et al., 2008b; Seidl et al., 2006), represses capsule formation, and activates biofilm formation in a glucose-rich environment (Seidl et al., 2008a). Indeed a *S. aureus ccp*A deletion mutant strain was less pathogenic than the wild-type strain in a murine abscess model (Li et al., 2010) and why *ccpA* inactivation increased the susceptibility of hyperglycemic animals to acute pneumonia infections (Bischoff et al., 2017). Nevertheless, the mechanism by which CcpA affected *S. aureus* pathogenesis cannot be simply resumed as a modulation of the RNAIII-dependent regulatory networks. Therefore, it has been suggested that CcpA can also act indirectly on gene expression through the action of other regulatory proteins or sRNAs (Somerville and Proctor, 2009; Richardson et al., 2015).

In Enterobacteriaceae, several small non-coding RNAs (sRNAs) have been shown as key actors of the uptake and the metabolism of carbohydrates (reviewed in Bobrovskyy and Vanderpool, 2014). For instance, they participate in the regulation of the galactose operon and carbon catabolite repression, metabolism of amino acids, and contribute to bacterial survival during phospho-sugar stress. The importance of sRNAs in regulatory networks is now well recognized to rapidly adjust cell growth to various stresses and changes in the environment. Thus, they are obvious candidates creating the links between virulence and metabolism. One example is RsaE, a sRNA conserved in *S. aureus* and *Bacillus subtilis*, that controls enzymes involved in the TCA cycle (Geissmann et al., 2009; Bohn et al., 2010). Recent observations showed that *B. subtilis* ResD represses RoxS (homologous to *S. aureus* RsaE) to readjust the pool of NAD+/NADH in responses to various stress and stimuli (Durand et al., 2017). Its promoter is also highly conserved among *Staphylococceae* and is recognized by the orthologous response regulator SrrA in *S. aureus*. Responding to reactive oxygen species through SrrAB, *S. aureus* RsaE may also intervene in the survival of cells against host immune reactions (Durand et al., 2015; Durand et al., 2017).

Here, we have identified a signaling pathway responding to glucose uptake, which involves a sRNA, called RsaI. This 144 nucleotides-long sRNA is highly conserved among *Staphylococcacea*, and carries two conserved regions including two G-track sequences and a long unpaired interhelical region rich in pyrimidines (Geissmann et al., 2009; Marchais et al., 2010). RsaI is strongly expressed at the stationary phase of growth in rich medium (Geissmann et al., 2009), and is enhanced after vancomycin exposure (Howden et al., 2013). In this study, we revealed that CcpA is the main repressor of RsaI expression in the presence of glucose, and that this inhibition is alleviated after the utilization of glucose. The identification of the targetome of RsaI using the MS2-affinity purification approach coupled to RNA sequencing (MAPS), unexpectedly showed two classes of RNA targets, including mRNAs involved in glucose uptake, sugar metabolism, and biofilm formation, as well as various sRNAs. Using site-directed mutagenesis, we identified two functional regions of RsaI with distinct regulatory functions.

Taken together, these data illustrate how a multifunctional RNA provides a signal for a metabolic switch when the preferred carbon source is metabolized, and may serve as an RNA sponge to control the balance between two essential pathways responding to either glucose or glucose 6-phosphate uptake. We will discuss the importance of sRNA-mediated regulation in *S. aureus* to fine tune the expression of genes according to essential nutrient availability, and their consequences on metabolism adaptation and virulence.

## RESULTS

### The expression of RsaI is inhibited by glucose and by the carbon catabolite protein A

We have previously shown that the synthesis of RsaI is high at the stationary phase of growth in the BHI medium while its expression was considerably enhanced at the exponential phase of growth in MHB medium (Geissmann et al., 2009). These data suggested that RsaI expression is regulated in a manner dependent on nutrient or biosynthetic intermediate availability. An obvious difference between BHI and MHB composition is their carbon source, glucose and starch, respectively. We wondered if the expression of RsaI might be dependent on the available carbon source. For this purpose, Northern blot experiments were performed on total RNAs prepared from HG001 (wild type) strain grown in the MHB medium at various time points (Figure 1A, 1B). In MHB where the glucose is not immediately available as the carbon source, RsaI was constitutively and highly expressed (Figure 1A). Conversely, when glucose was added, either at the beginning of the culture or after 3 h of growth, transcription of RsaI was immediately stopped, indicating the repressing effect of this sugar (Figure 1A).

**Figure 1:**
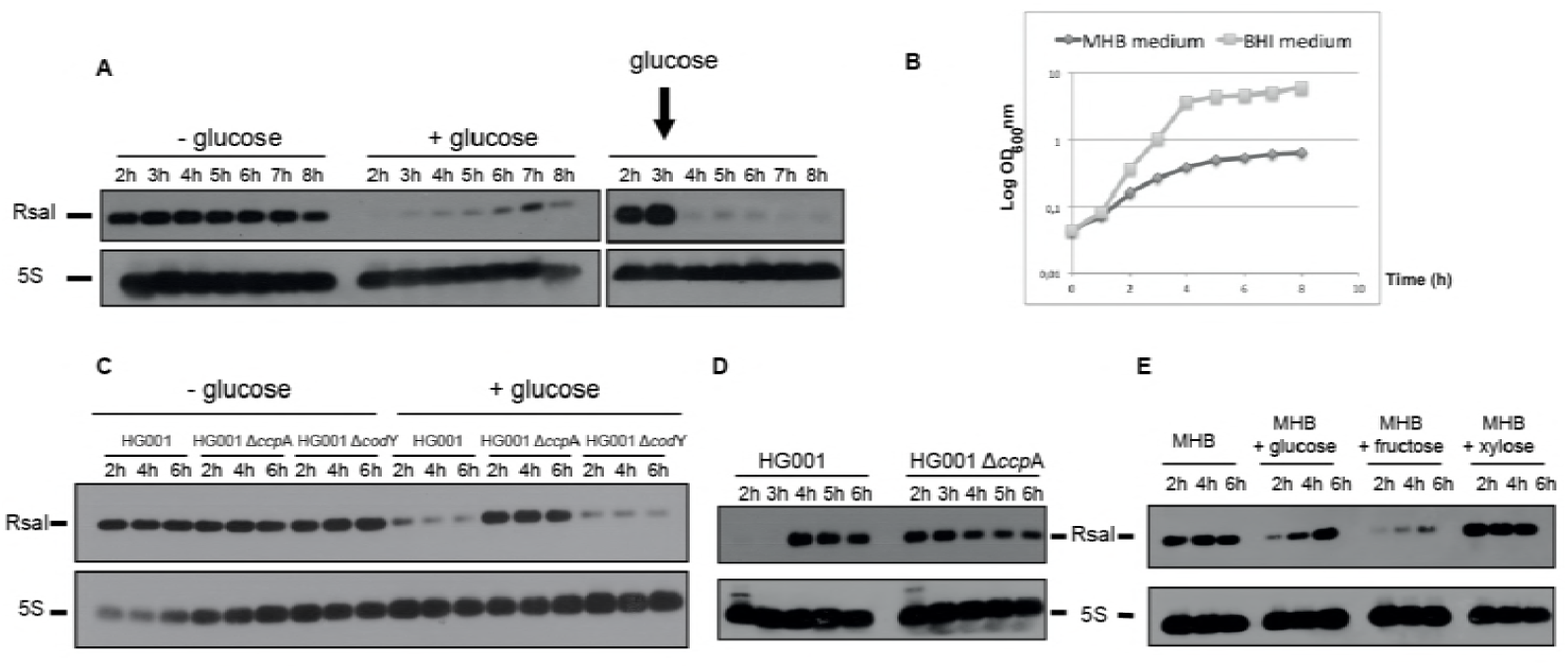
RsaI responds to glucose through the transcriptional factor CcpA. **(A)** Northern blot experiments show the expression of RsaI during growth phase in the HG001 strain in MHB medium with or without the addition of 1,5 g/L of D-glucose at the beginning of growth (middle panel) or after 3 h of growth (right panel). **(B)** Growth curves of HG001 in MHB or BHI. **(C)** Northern blot experiment shows the expression of RsaI during growth phase in the HG001, Δ*ccp*A mutant strain, and Δ*cod*Y mutant strain, in MHB medium with or without the addition of 1,5 g/L of D-glucose. **(D)** Northern blot experiment shows the expression of RsaI in the HG001 and Δ*ccp*A mutant strains. Total RNA was prepared after 2, 3, 4, 5 and 6 h of culture in BHI medium at 37°C. **(E)** Northern blot analysis of RsaI in MHB medium with or without the addition of 1,5 g/L of glucose, fructose or xylose. For all the experiments shown in panels A and C to E, loading controls were done based on the expression of 5S rRNA (5S) revealed after hybridization of the membranes with a specific oligonucleotide. However for these controls, we used aliquots of the same RNA preparations but the migration of RNA samples was performed in parallel to the experiments on a separate agarose gel.

As CcpA sensed the intracellular concentration of glucose (Seidl et al. 2006), we analyzed if the expression of RsaI was CcpA-dependent. Northern blot analysis was performed on total RNA extracts prepared from HG001 strain and an isogenic mutant deleted of *ccpA* gene (Δ*ccpA*). The same experiment was performed with a mutant strain lacking the *codY* gene, which was shown as a direct regulator of amino acid biosynthesis, transport of macromolecules, and virulence (Majerczyk et al., 2010; Pohl et al., 2009). In the absence of glucose, the yield of RsaI was similar in all strains. However, in the MHB medium supplemented with glucose, the expression of RsaI dropped dramatically in the WT and Δ*codY* strains, whereas the expression of RsaI was still high in Δ*ccp*A strain (Figure 1C). In BHI medium, RsaI was expressed after 4 h of growth, while its expression was constitutive in the mutant Δ*ccpA* strain (Figures 1D, S1A).

We also analyzed the expression of RsaI in MHB medium supplemented with various sugars such as fructose and xylose (Figure 1E). The data showed that the expression of RsaI is very low at the beginning of growth in MHB supplemented with either glucose or fructose while the expression of RsaI is constitutive in MHB supplemented with xylose. These data suggest that the inhibition of RsaI transcription is only dependent of hexoses (Figure 1E).

Overall, we showed that CcpA represses RsaI expression in the presence of glucose. In accordance with this data, a conserved *cre* (GGAAAcGcTTACAT) sequence was found at position -30 upstream the transcriptional start site of RsaI (Figure S1B). This region was sufficient to confer repression by glucose in the complemented strain containing pCN51::P*rsa*I (data not shown).

### The targetome of RsaI as revealed by the MAPS approach

The MAPS approach (“MS2 affinity purification coupled to RNA sequencing”) was used to purify *in vivo* regulatory complexes involving RsaI. MAPS has been successful to identify the RNA targets of any sRNAs in *E. coli* (Lalaouna et al., 2015), and more recently of RsaA sRNA in *S. aureus* (Tomasini et al., 2017). Briefly, the MS2 tagged version of RsaI was expressed from a plasmid under the control of the *agr*-dependent P3 promoter, allowing an accumulation of the RNA at the stationary phase of growth in the Δ*rsaI* mutant strain. RsaI was detected by Northern blot using total RNAs extracted at 2, 4 and 6 h of growth in BHI medium. Using a DIG-labeled RsaI probe, we showed that the steady state levels of MS2-RsaI were very similar to the wild type (WT) RsaI, and that MS2-RsaI was specifically retained by the MS2-MBP fusion protein after the affinity chromatography (Figure S1C). The RNAs were then extracted from the eluted fraction to be sequenced. The data were analyzed using Galaxy (Afgan et al., 2016) and the sequencing reads were mapped with the reference HG001 genome (Caldelari et al., 2017; Herbert et al., 2010) counted per feature and normalized. The enrichment of putative targets was done by comparing the number of reads obtained from the MS2-RsaI purification and the MS2 alone as control. The data were reproduced in two independent experiments (Tables 1, S3).

**Table 1:**
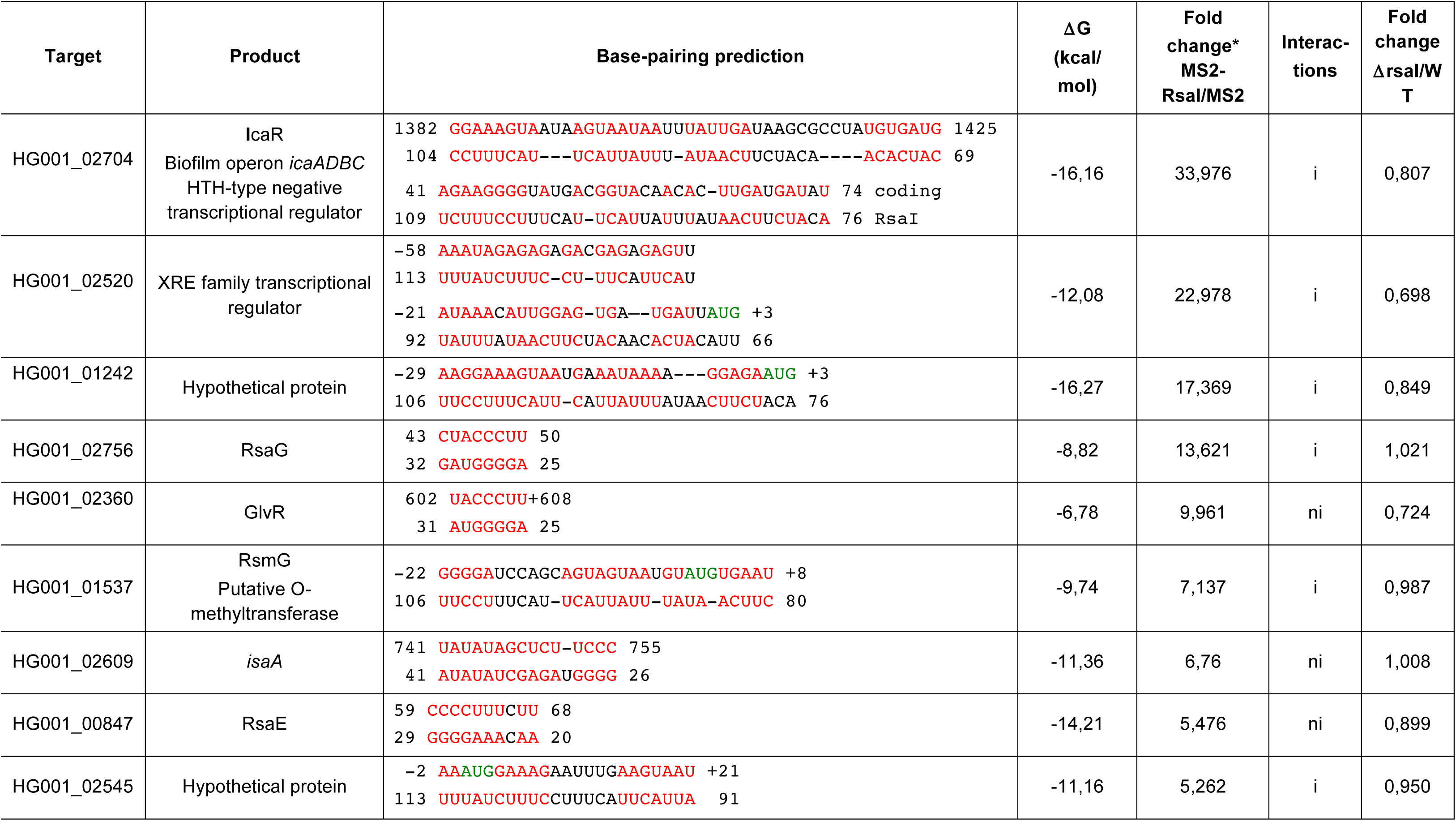
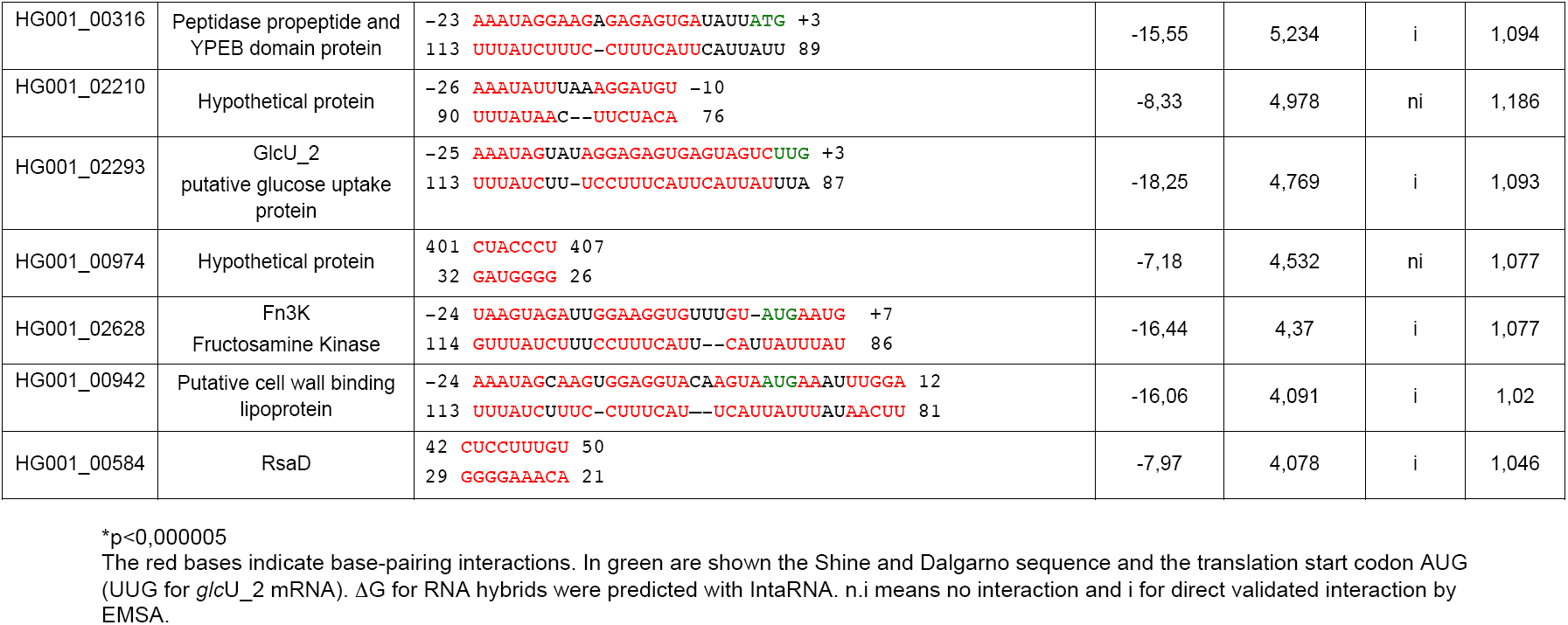
*In silico* predictions of base-pairing interactions for selected RNAs found by MAPS with MS2-RsaI.

Interestingly, two classes of RNAs were co-purified with RsaI, including several mRNAs and sRNAs. The best mRNA candidate encodes IcaR, the repressor of the *icaADBC* operon, which is required for the synthesis of the poly-b-1,6-N-acetylglucosamine polymer (PIA-PNAG), the main staphylococcal exopolysaccharide in biofilm. Several mRNAs encode proteins linked to sugar utilization and transport, such as *glcU*_2 encoding a major transporter of glucose, and *fn3K* encoding fructosamine 3-kinase, and other less enriched mRNAs (< 4-fold) produce the trehalose-specific PTS transporter (TreB), a sugar phosphatase (YidA), and a maltose transport system permease (Table S3). Finally, several mRNAs express transcriptional regulators (Xre type, the maltose regulatory protein GlvR, SlyA, SigS). As for sRNAs, we found RsaD, RsaE, RsaG, and the less enriched RsaH, which all contained at least one conserved C-rich motif (Geissmann et al., 2009).

We then searched for possible intermolecular base-pairing interactions between RsaI and its potential RNA targets using IntaRNA (Table 1). Stable interactions were predicted for most of the enriched RNAs. The CU-rich unpaired region of RsaI was predicted to form base-pairings with most of the mRNAs. They were located close or at the ribosome binding sites of most of the mRNAs except for *icaR* and *isaA*, which involves nucleotides in their 3’ untranslated region (Table 1). A second domain of interaction corresponded to the G-track sequences located in the first hairpin domain of RsaI and the C-rich sequences of the sRNAs RsaD, RsaE, RsaG and RsaH.

These data suggested that RsaI is a multifunctional RNA, which regulates the expression of mRNAs at the post-transcriptional level, while its activities might be regulated by other sRNAs.

### RsaI contains two distinct regulatory domains

Based on the MAPS data, we first analyzed whether RsaI effectively binds to the mRNA and sRNA candidates using gel retardation assays (Figures 2 B, C and S2). *In vitro* 5’ end-labeled RsaI was incubated with increasing concentrations of various mRNAs, for example, those encoding proteins involved in biofilm formation (IcaR), sugar uptake and metabolism (GlvR, GlcU_2, IsaA, and Fn3K), and several sRNAs (RsaG, RsaE, and RsaD). For these experiments, we used the full-length mRNAs and sRNAs (Table S2). Complex formation was performed with RNAs, which were renatured separately in a buffer containing magnesium and salt. As expected, the data showed that RsaI formed complexes with high affinity (between 20-100 nM, Table 1) with many RNAs such as *ica*R, *glc*U_2, and *fn*3K mRNAs, and RsaG sRNA (Figure 2). The stability of other complexes (e.g. *tre*B mRNA and RsaD sRNA) was significantly lowered (> 250 nM) (Figure S2).

**Figure 2:**
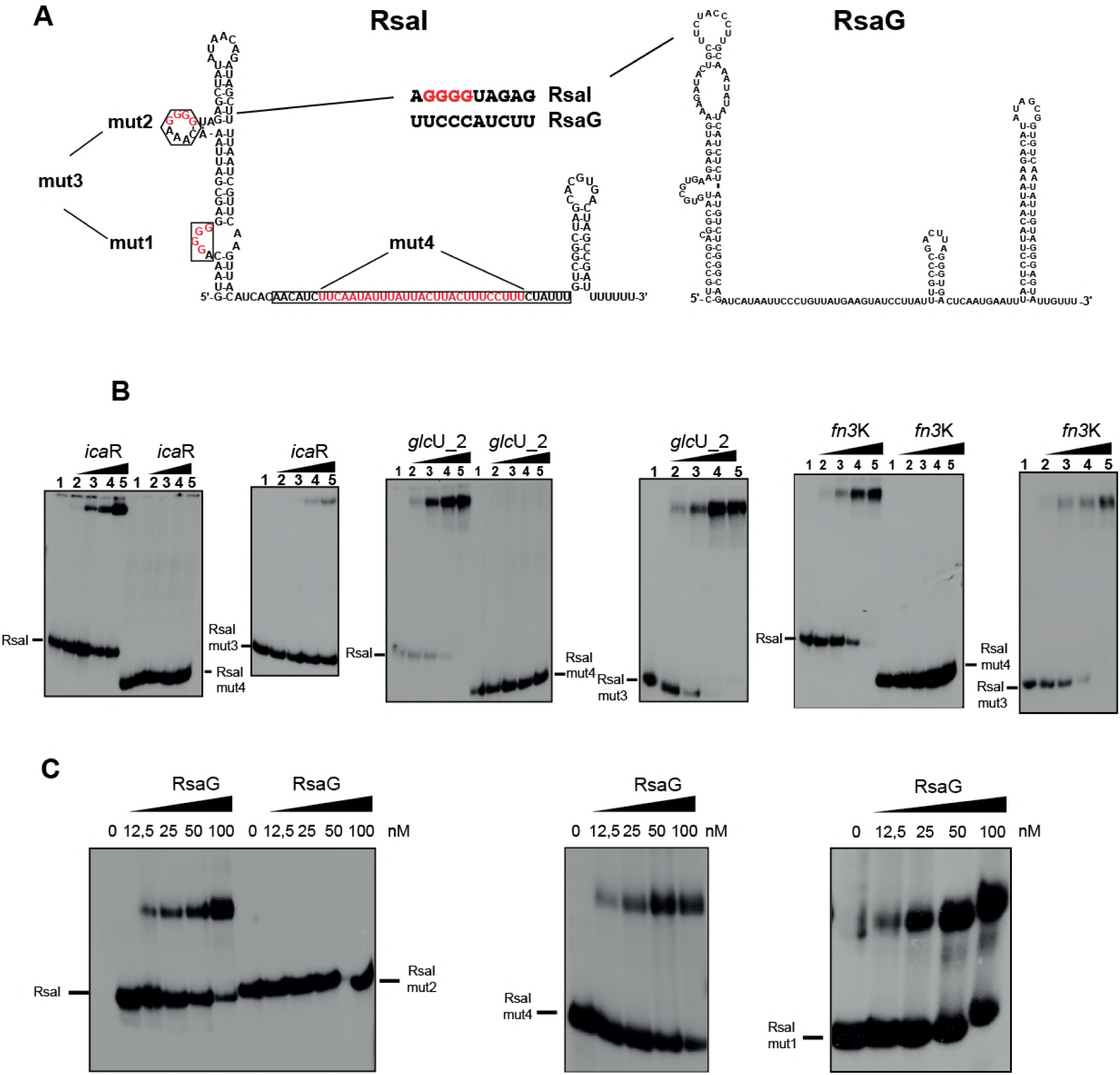
RsaI binds to *ica*R, *glc*U_2, *fn3*K mRNA and the sRNA RsaG. **(A)** Secondary structure model of RsaI. In red, are the nucleotides deleted in the RsaI mutants (mut1 to mut4). The potential base-pairings between RsaI and RsaG are shown. Squared nucleotides are conserved sequences in RsaI. **(B)** Gel retardation assays show the formation of the complex between RsaI and *ica*R, *glc*U_2 and *fn*3K. The 5’ end-labeled wild-type RsaI (RsaI), RsaI mutant 3 (RsaI mut3), and RsaI mutant 4 (RsaI mut4) were incubated with increasing concentration of mRNAs: lane 1, 10 nM; lane 2, 20 nM; lane 3, 50 nM; lane 4, 100 nM and lane 5, 200 nM. **(C)** Gel retardation assays show the formation of the complex between RsaI and RsaG. The 5’ end-labeled wild-type RsaI (RsaI), RsaI mutant 2 (RsaI mut2), RsaI mutant 4 (RsaI mut4), and RsaI mutant 1 (RsaI mut1) were incubated with increasing concentrations of RsaG given in nM on the top of the autoradiographies.

Based on the base-pairing predictions, mutations have been introduced into *rsa*I to map its functional regulatory regions. The two G-track sequences were mutated separately (RsaI mut1: ΔG7-G10, RsaI mut2: ΔG26-G29) or together (RsaI mut3: ΔG7-G10/ΔG26-G29), while several nucleotides (RsaI mut4: ΔU81-U107) were deleted in the interhelical unpaired sequence (Figure 2A). We then analyzed the ability of mutated RsaI derivatives to form complexes with *glc*U_2, *fn*3K, and *ica*R mRNAs, and RsaG sRNA (Figure 2B). The data showed that RsaI mut3 bind to all three mRNAs similarly to the WT RsaI, while complex formation was completely abolished with RsaI mut4. Only the mutations in the second G-track sequences (RsaI mut 2) strongly altered the binding to RsaG, while the two other mutated RsaI (mut1 and mut4) recognized RsaG with an equivalent binding affinity as the WT RsaI (Figure 2C).

The MAPS approach was also used to monitor the effect of the mutations in RsaI on its target RNAs *in vivo*. The MS2 tagged mutant versions of RsaI (mut2 and mut4) were expressed in the HG001Δ*rsaI* mutant strain. As described above, the enrichment of putative targets was calculated by comparing the number of reads obtained from MS2-RsaI purification and the MS2 alone as control (Table S4). In the fraction containing MS2-RsaI mut4, the mRNA targets encoding IcaR, Fn3K, and GlcU_2 were strongly reduced while RsaG was still significantly enriched. Conversely, we observed that the three mRNAs were still co-purified together with the MS2-RsaI mut2 at a significant level close to the WT RsaI while RsaG was strongly reduced in the fraction containing MS2-RsaI mut2.

The MAPS experiments are well correlated with the gel retardation assays. Hence, they revealed that RsaI has at least two distinct regulatory domains that directly interact either with mRNAs or with sRNAs.

### The effect of RsaI on global gene transcription in *S. aureus*

In order to get a global overview of RsaI impact on gene regulation, a comparative transcriptomic analysis was performed on total RNAs extracted from the WT HG001 strain, the isogenic HG001Δ*rsa*I mutant strain, and the same mutant strain complemented with a plasmid expressing RsaI under the control of its own promoter (Table S5). The cultures were done in triplicates with high reproducibility in BHI medium at 37°C until 6h, under the conditions allowing the expression of RsaI (Figure 1). We have considered a gene to be regulated by RsaI if the ratio between two strains is at least twofold. Significant differences were mostly observed between the mutant Δ*rsa*I versus the same strain expressing RsaI from a plasmid. Accordingly, the mRNA levels of 26 and 50 CDS were significantly decreased and enhanced, respectively, when the complemented strain was compared to the mutant Δ*rsa*I (Table S5). Most of the RsaI-dependent activation was observed for genes involved in fermentation processes (i.e., *ldh*1 encoding lactate dehydrogenase, *adh*, encoding alcool dehydrogenase, *pfl*B encoding formate acetyltransferase, *foc*A encoding one of the formate transporter, and *pfl*A encoding pyruvate-formate lyase activating enzyme), in energy-generating processing (i.e., *qox*ABCD operon encoding terminal oxidases for aerobic respiration, *cta*A encoding heme synthase, *hmp* encoding flavoprotein), in amino acid metabolism (i.e., *arg* encoding arginase, *pro*P encoding proline/betaine transporter, *tdc*B encoding L-threonine dehydratase catabolic), in peptide transport system (*opp*B/D), and in the biosynthesis of co-factors and prosynthetic groups (i.e., *nas*E*-nas*F operon). The levels of two sRNAs (RsaG, RsaH) were also enhanced in the HG001 WT strain. Besides, the overexpression of RsaI caused a reduced expression of genes that are functionally related. Several of them are involved in glycolysis and pentose phosphate pathway (i.e., *fba*1 encoding fructose bi-phosphate aldolase, *gnd* encoding 6-phosphogluconate dehydrogenase, *tkt* encoding transkelotase, *tre*R_2 encoding transcriptional repressor of the trehalose operon containing *tre*B), in thiamine co-factor synthesis (*thi*DME operon, *ten*A encoding a putative thiaminase, *thi*W encoding a thiamine transporter), in carbohydrate uptake (*pts*I encoding phosphoenolpyruvate-protein phosphotransferase), and in arginine catabolism (*arc* operon). Additionally, significant repression was also observed for *mia*B encoding tRNA specific modification enzyme, *tyr*S encoding tryptophanyl-tRNA synthetase, and *ebp*S encoding the cell surface elastin binding protein. Interestingly, the MAPS approach revealed that several of these mRNAs (i.e., *qox*ABCD operon, *tyr*S, *arc*C1, *tre*R_2, *plf*A*-plf*B) were enriched together with RsaI (Table S3). In contrast, most of RsaI targets identified by MAPS did not show significant mRNA level variations when RsaI was deleted or overexpressed suggesting RsaI regulates the translation of these mRNAs.

These data showed that high concentrations of RsaI affected the mRNA levels of several enzymes involved in sugar metabolism, in the pentose phosphate pathway, and various processes linked to energy production.

### RsaI inhibits the translation of several mRNAs by masking their RBS

We then analyzed whether RsaI would preferentially regulate translation of *glc*U_2 and *fn*3K mRNA targets because their levels were not altered in the Δ*rsaI* mutant strain and the C/U unpaired region of RsaI was predicted to base-pair with their SD (Figure 2). Using toe-printing assays, we analyzed the effect of RsaI on the formation of the ternary initiation complex constituting of mRNA, the initiator tRNA, and the 30S subunit. For both mRNAs, the formation of the ternary initiation complex is illustrated by the presence of a toe-print signal at position +16 (+1 being the initiation codon). For *fn*3K mRNA, two toe-print signals were observed at +16 and at +20, most probably corresponding to the presence of two AUG codons distant of 5 nucleotides. However, only the first AUG is used *in vivo* (Gemayel et al., 2007). The addition of increasing concentrations of RsaI together with the 30S strongly decreased the toe-print signals for both mRNAs showing that RsaI is able to form a stable complex with the mRNA sufficient to prevent the binding of the 30S subunits (Figure 3A).

**Figure 3:**
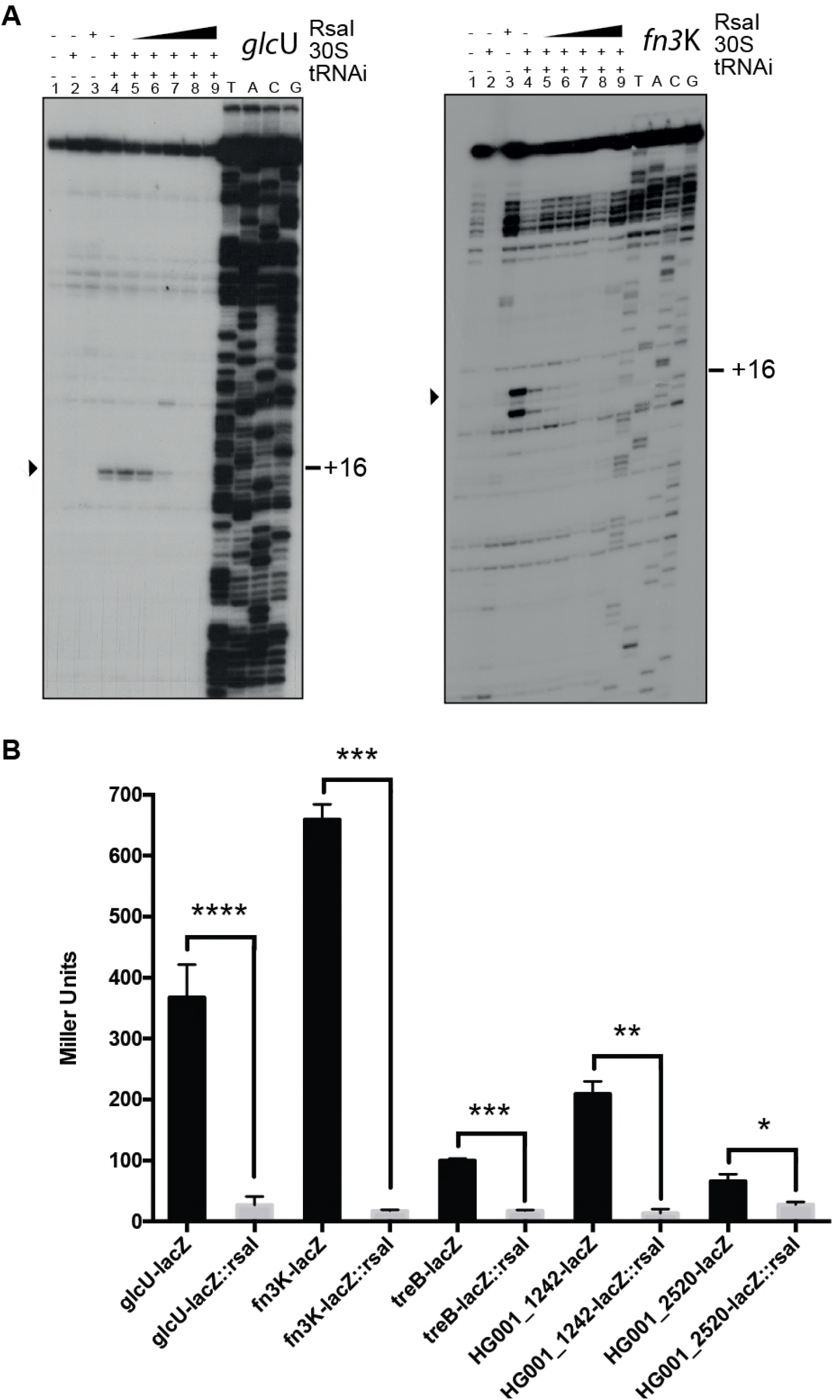
RsaI inhibits *glc*U_2 and *fn*3K translation. **(A)** Toe-print assay showing the effect of RsaI on the formation of the ribosomal initiation complex of *glc*U_2 and *fn*3K respectively. Lane 1 : incubation control of mRNA; lane 2 : incubation control of mRNA with 30S subunits; lane 3 : formation of the ribosomal initiation complex containing mRNA, 30S and the initiator tRNA^fMet^ (tRNAi); lane 4: incubation control of mRNA with RsaI; lane 5 to 9 : formation of the initiation complex in the presence of increasing concentrations of RsaI, respectively : 50, 100, 150, 300 and 400 nM. Lanes U, A, C, G: sequencing ladders. The Shine and Dalgarno (SD) sequence, the start site of translation (A +1 of the AUG initiation codon) and the toe-printing signals (N+16) are indicated. **(B)** Theç -galactosidase activity (Miller Units) have been measured from P*rpo*B::*glc*U::*lac*Z::P*rsa*I, P*rpo*B::*fn*3K::*lac*Z::P*rsa*I, P*rpo*B::*tre*B::*lac*Z::P*rsa*I and P*rpo*B::*HG001_2520*::*lac*Z::P*rsa*I expressed in the mutant strain HG001-Δ*rsa*I. Theç -galactosidase activity was normalized for bacterial density and the results represented the mean of four independent experiments. * p < 0.05, ** p < 0.005,*** p < 0.0005, **** p < 0.0001.

To further validate the *in vivo* relevance of RsaI-dependent repression of the mRNA *glc*U_2, *fn*3*K, tre*B, *HG001_01242,* we analyzed the expression of a reporter construct carrying their regulatory regions fused to *lac*Z and expressing RsaI under its own promoter in *S. aureus* HG001Δ*rsaI* strains. The ß-galactosidase activity was reproducibly found higher in the Δ*rsa*I mutant strain for all reporter constructs (Figure 3B) showing that the absence of RsaI alleviated the repression of the reporter gene.

These data strongly suggested that RsaI primarily regulated translation initiation of *glc*U_2, *fn*3K, *tre*B, *HG001_01242* and *HG001_02520* mRNAs mainly by masking their ribosome binding sites.

### RsaI interacts with *ica*R mRNA and affects PIA-PNAG synthesis

We have shown that RsaI is able to form stable base-pairings with *ica*R mRNA that encodes the repressor of the main exopolysaccharidic compound of *S. aureus* biofilm matrix. This mRNA is of particular interest because it contains a large 3’ UTR that is able to bind to its own Shine and Dalgarno (SD) sequence (Ruiz de los Mozos et al., 2013). Consequently, the long-range interaction provokes an inhibitory effect on translation and generates a new cleavage site for RNase III (Ruiz de los Mozos et al., 2013). RsaI is predicted to form base-pairings with the 3’UTR of *ica*R downstream of the anti-SD sequence (Table 1). In a first experiment, we monitored whether the long-range interaction might be critical for RsaI binding. Previous work showed that the substitution of UCCCCUG sequence by AGGGGAC, located in the 3’UTR of *ica*R, significantly destabilized the long-range interaction to enhance *ica*R translation (Ruiz de los Mozos et al., 2013). However, gel retardation assays showed that the WT and the mutant *ica*R mRNAs bind to RsaI with an equivalent binding affinity (Figure 2B), suggesting that the anti-SD region is not required for RsaI binding. Instead, we have shown that the 3’UTR of *icaR* contained the high affinity binding site for RsaI (Figure 4A, Table 1).

**Figure 4:**
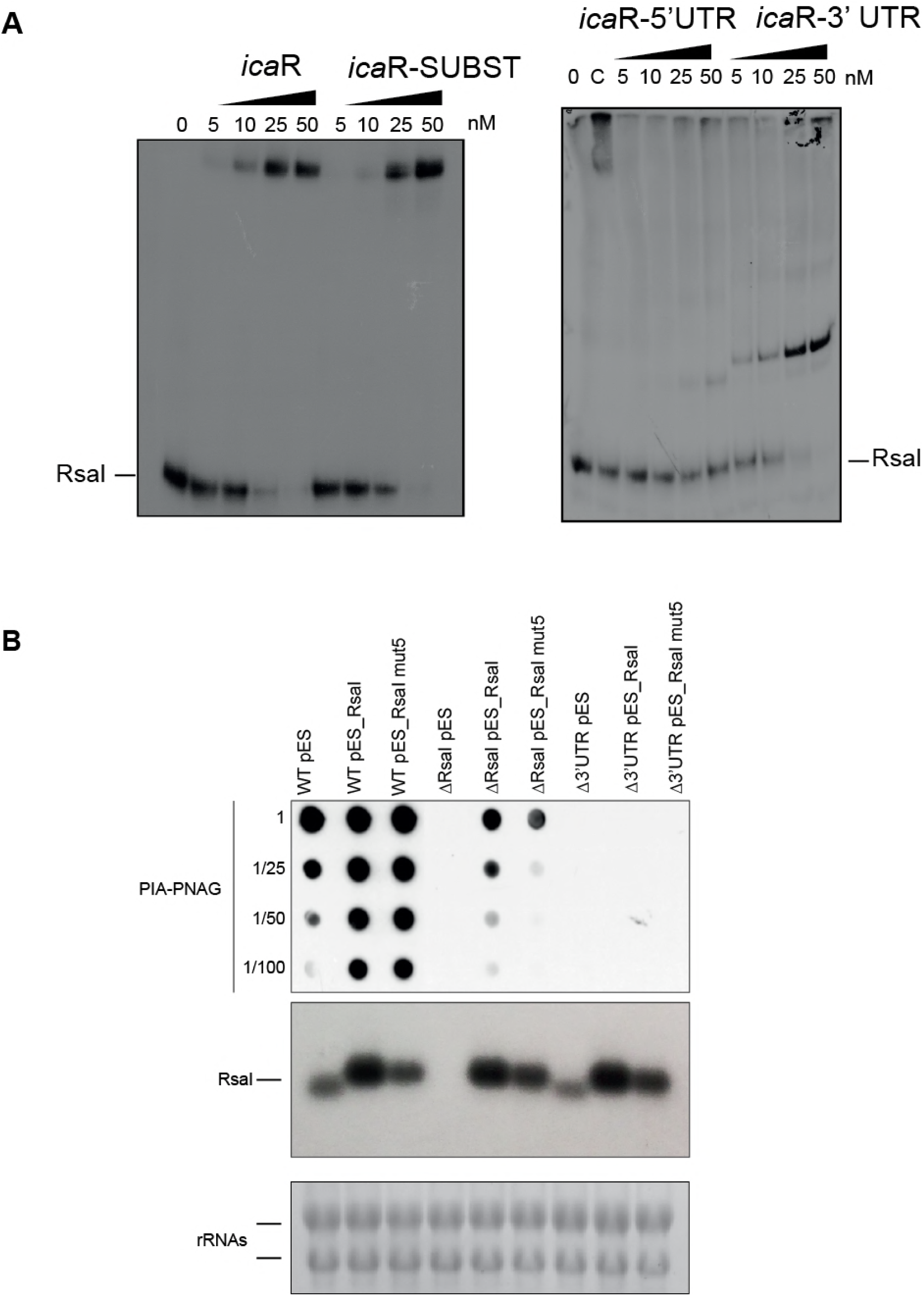
Interaction of RsaI to *ica*R mRNA induces biofilm production. (**A**) Gel retardation assays show the formation of the complex between RsaI and *ica*R full length, *ica*R SUBST, *ica*R-5’ UTR, and *ica*R-3’ UTR. The 5’ end-labeled of RsaI was incubated with increasing concentrations of mRNAs. UTR is for untranslated region. SUBST stands for the substitution of UCCCCUG sequence by AGGGGAC, located in the 3’UTR of *ica*R significantly destabilizing the long-range interaction to enhance *ica*R translation **(B)** *In vivo* effects of RsaI on PIA-PNAG synthesis in the *S. aureus* wild-type (WT) 132 strain, the Δ*rsa*I mutant and the strain with a deletion of *ica*R 3’ UTR (Δ3’UTR) containing the pES plasmids with *rsa*I and *rsa*I mut5. Quantification of PIA-PNAG exopolysaccharide biosynthesis using dot-blot assays. Serial dilutions (1/5) of the samples were spotted onto nitrocellulose membranes and PIA-PNAG production was detected with specific anti-PIA-PNAG antibodies. RsaI was detected in the same samples by Northern blot using a probe directed against RsaI. Ethidium bromide staining of rRNA was used as loading control.

We then analyzed the capacity of the WT and mutant Δ*rsa*I strains to synthesize PIA-PNAG exopolysaccharides. Dot-blot assays were performed with anti PIA-PNAG specific antibodies to monitor the levels of PIA-PNAG in the strain 132, which produced high levels of this exopolysaccharide in the presence of NaCl. We tested the PIA-PNAG levels in WT strain, the isogenic Δ*rsa*I mutant strain and the Δ3’UTR *ica*R mutant strain carrying a deletion of the 3’UTR of *ica*R grown for 6 h in TSB containing NaCl (Figure 4B). The data suggested that RsaI and the 3’UTR of *ica*R are required for efficient production of PIA-PNAG because only the WT strain produces significant levels of exopolysaccharides. The three strains were also transformed with a plasmid overexpressing RsaI or RsaI mut5 carrying a substitution of nucleotides 88 to 103 (TTATTACTTACTTTCC to AATAATGAATGAAAGG). This mutation is expected to decrease the stability of RsaI-*ica*R duplex. Northern blots were performed to control that RsaI was properly over-expressed in these strains (Figure 4B). In the WT background, which expresses the endogenous RsaI, the additional overexpression of WT or the mutated version of RsaI causes a similar enhanced synthesis of PIA-PNAG. However, in the mutant Δ*rsa*I strain background, the expression of RsaI mut5 induces the accumulation of lower levels of PIA-PNAG than the strain expressing the WT RsaI (Figure 4B). Finally, the synthesis of PIA-PNAG is completely inhibited in the three Δ3’UTR mutant strains indicating that in this background, RsaI is not able to exert its regulatory functions.

Taken together, these data suggest that RsaI would control PIA-PNAG synthesis by reducing the IcaR repressor protein levels through a specific interaction with the 3’UTR of *ica*R mRNA.

### RsaI connects sugar metabolism and NO stress responses through sRNA binding

Three sRNAs (RsaD, RsaE, and RsaG) were significantly enriched together with RsaI. RsaG is the most enriched sRNA, and the complex formed between RsaI and RsaG was highly stable (Figure 2). We first addressed the consequences of such pairings on sRNA regulation. Using gel retardation assays, we analyzed whether RsaG is able to form a ternary complex with RsaI and one of its target mRNA, or if RsaG competes with the mRNA for RsaI binding. In this experiment, we used a defined concentration of RsaG (50 nM) known to be sufficient to bind most of RsaI molecules, which were 5’ end labeled. In the absence of RsaG, a concentration dependence of the mRNA *glc*U_2 shows that it binds efficiently to RsaI. The addition of RsaG induces the formation of a high molecular weight complex formed by RsaG, RsaI and the mRNA (Figure 5A). The same results were obtained with other mRNA targets of RsaI (*HG001_1242, HG001_0942*) (Figure 5A). The formation of a ternary complex is a good indication that RsaG binding does not interfere with the regulatory functions of RsaI. In addition, using rifampicin treatment, we have controlled that the half-lives of RsaI or RsaG were not significantly modified in the absence of RsaG or RsaI, respectively (Figure 5B). Thus *in vivo*, the absence of one sRNA did not impact transcription or stability of the other.

We have previously shown that RsaG is expressed at the stationary phase of growth in BHI medium (Geissmann et al, 2009). Interestingly, *rsa*G gene is localized just downstream the mRNA encoding the hexose phosphate transporter UhpT, whose transcription is activated by the two component system HptRS in response to extracellular glucose-6 phosphate (G-6P), another important carbon source produced by host cells (Figure S3A, Park et al, 2015). We therefore analyzed whether RsaG expression would also respond to the cellular concentration of G-6P. Northern blot experiments were performed on total RNA extracts produced from the WT HG001 strain grown in BHI medium supplemented with G-6P. Under these conditions, the synthesis of RsaG appears to be strongly enhanced (Figure S3B, left panel). The deletion of *hpt*RS considerably reduced the levels of RsaG. Therefore, these data strongly indicate that RsaG is activated by HptRS upon G-6P signaling together with *uhp*T (Figure S3B, right panel) and suggest that RsaG sRNA might be derived from the *uhp*T 3’UTR.

**Figure 5:**
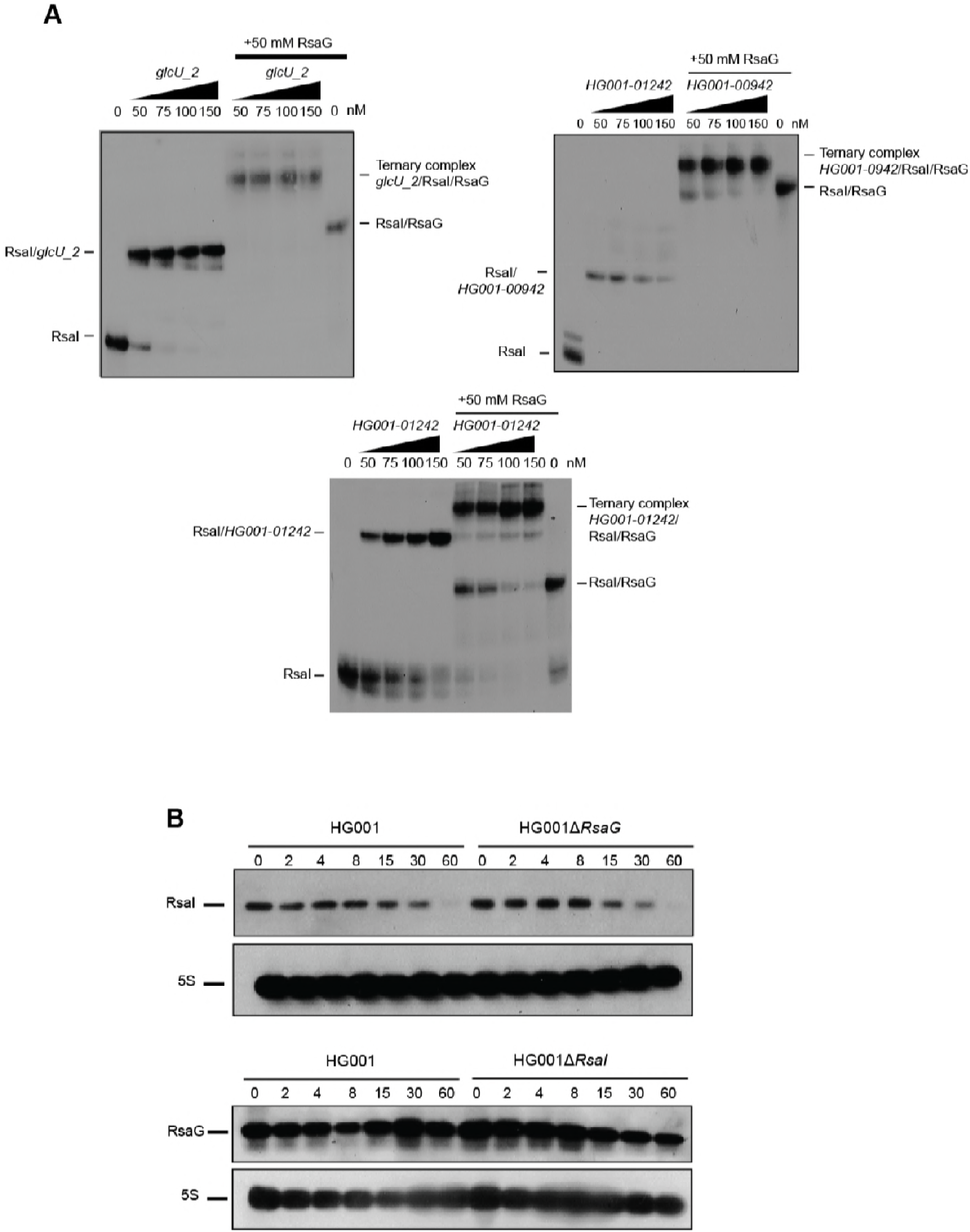
Regulatory function of RsaI is not impaired by binding of RsaG. **(A)** Ternary complex formation between RsaI, mRNA target *glc*U_2, *HG001_00942 and HG001_1242*, and RsaG. The 5’-end labeled RsaI was incubated with increasing concentrations of *glcU_2* alone, or in the presence of 50 nM of RsaG**. (B)** Northern blots experiments have been used to monitor the half-lives of RsaI and RsaG in either HG001-Δ*rsa*G or HG001-Δ*rsa*I mutant strains. The cells were treated with rifampicin at 4h of growth and total RNAs were extracted after 2, 4, 8, 15, 30 and 60 minutes at 37°C in BHI. 5S rRNA was probed to quantify the yield of RNA in each lane using the same samples but run on two different gels.

We also analyzed the signaling pathway of RsaD. This sRNA was previously shown to accumulate at the stationary phase of growth and its expression was enhanced under heat shock conditions (Geissmann et al., 2009). We then demonstrated that the expression of RsaD is not dependent on the two-component system SaeRS but is strongly enhanced by SrrAB. Indeed, its expression drops considerably in a Δ*srr*AB mutant strain (Figure S3C). Because SrrAB is able to sense and respond to nitric oxide (NO) and hypoxia (Kinkel et al., 2013), we tested the effect of NO on RsaD synthesis by adding diethylamine-NONOate, as previously described (Durand et al., 2015). We observed a significant and reproducible increase in RsaD expression in the WT strain about 10 min after the addition of diethylamine-NONOate to the medium (Figure S3D). RsaD expression decreased after 20 min due to the short half-life of diethylamine-NONOate. Upstream *rsa*D, we identified a conserved motif AGTGACAA that could be responsible for the SrrAB-dependent transcription. It is of interest that the synthesis of RsaE was also shown to be under the control of SrrAB (Durand et al., 2015).

These data suggested that through the binding of sRNAs, RsaI would link sugar metabolism pathways, carbon source utilization, energy production, and stress responses.

## DISCUSSION

In this work, we have investigated the cellular functions of one abundant sRNA, called RsaI (or RsaOG), which is highly conserved among *Staphylococcaea* (Geissmann et al., 2009; Marchais et al., 2010). In contrast to many sRNAs that contained C-track motifs, this RNA has the particularity to carry two conserved G-rich sequences and a large unpaired CU-rich region (Figure 1). This RNA was proposed to fold into a pseudoknot structure involving the two conserved regions limiting the access of regulatory regions (Marchais et al., 2010). Combining several global approaches (MAPS, transcriptomic analysis), we show that RsaI controls a large regulon involved in sugar uptake and metabolism, biosynthetic and co-factor synthesis, cytochrome biosynthesis, anaerobic metabolism, as well as iron-sulfur cluster repair, and NO detoxification (Figure 6). Additionally, RsaI emerges in *S. aureus* as a new class of regulatory RNAs acting as a sponge of several sRNAs carrying C-track sequences that are induced upon enhanced concentrations of specific metabolite (G-6P) or ROS signaling pathway (synthesis of NO). Hence, RsaI appears as a key regulator that links many adaptive pathways in response to the preferred carbon source availability.

**Figure 6:**
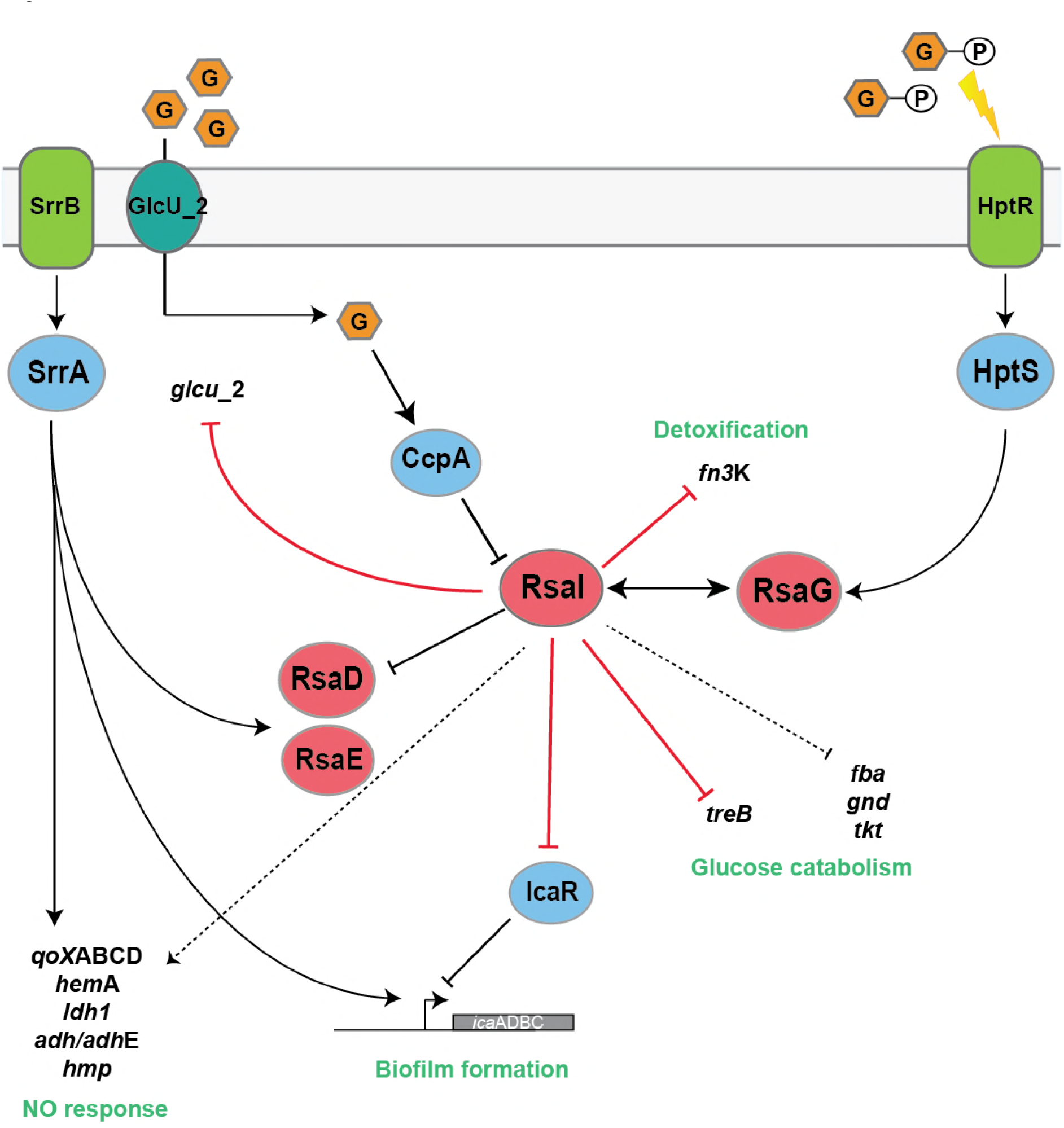
Schematic drawing summarizing the regulatory networks involving the bi-functional RsaI and its mRNA targets in *Staphylococcus aureus.* Arrows are for activation, bars for repression. In blue, are the transcriptional protein regulators, and in red the regulatory sRNAs. Red lines corresponded to post-transcriptional regulation and black lines to transcriptional regulation. Dotted line represented the regulatory events for which direct regulation is not yet demonstrated.

### The expression of RsaI is a signature of a metabolic switch responding to glucose

The cellular level of RsaI is tightly controlled during growth in rich medium. In this study, we showed that RsaI expression is strongly and rapidly repressed at the transcriptional level through the activity of CcpA in response to glucose availability. Carbon catabolite repression is a universal regulatory phenomenon that allows the cells to use the preferred carbon source to produce energy, and to provide the building blocks for macromolecules. Concomitantly it represses genes that are involved in the metabolism of less-preferred carbon sources. To do so, CcpA has to be activated through a cascade of events involving its co-regulator histidine-containing phosphocarrier protein (HPr), which has been phosphorylated by its cognate kinase/phosphorylase HprK/P activated in the presence of glycolytic intermediates. It is thought that binding of the phosphorylated HPr to CcpA enhances its DNA binding affinity to the *cre* binding site to repress or activate the target genes. RsaI repression requires the presence of CcpA, most probably through the binding at a *cre* motif located at -45/-30 (GGAAAACGCTTACAT) from the *rsa*I transcriptional start site (Figure 1). Interestingly, deletion of *ccp*A gene affected vancomycin resistance (Seidl et al., 2009) and recent observations showed that sub-inhibitory treatment of cells with vancomycin led to an enhanced expression of RsaI (Howden et al., 2013). We showed that the repression of RsaI is alleviated as soon as the concentration of glucose is strongly reduced. Therefore, RsaI might be a signature of a metabolic switch of the bacterial population. Using the MAPS approach, we could identify several mRNAs that strongly bind to RsaI with its long unpaired CU-rich region. Among these mRNAs, we show that *glc*U_2 encoding a glucose permease, *fn*3K encoding the fructosamine 3 kinase, and *tre*B encoding a kinase involved in trehalose transport are all regulating at the translational level by RsaI. This mechanism is most likely common to all mRNAs that could form base-pairings between their ribosome binding sites and the CU-rich region of RsaI. This is the case for mRNAs encoding a transcriptional regulator of the XRE family, the O-methyltransferase RsmG, a peptidase, a cell wall binding lipoprotein, and DegV containing protein (Table 1). Indeed, these mRNAs were no more found enriched together with RsaI_mut 4 in which the CU-rich region has been deleted (Table S3). Among these proteins, two of them are directly involved in sugar uptake and metabolism (GlcU_2, and TreB). In *S. aureus*, the PTS (phosphotransferase system)-dependent and independent transport systems ensure efficient glucose transport (reviewed in (Götz et al., 2006). If a rapidly metabolizable sugar (such as glucose) is used during growth at a rather low concentration, the transport will occur via the PTS system and concomitantly, the carbon catabolite repression system will be activated through CcpA protein. At high concentration of glucose, it is assumed that the sugar will be transported by the PTS system and in addition by the permease GlcU_2. The glucose transported by the permease will be phosphorylated within the cell by the glucose kinase GlkA. Hence, it is reasonable to propose that RsaI would repress the synthesis of GlcU_2 when the glucose concentration dropped. RsaI inhibits also key enzymes of the trehalose metabolism (TreB), a diholoside which is transported exclusively by PTS (Bassias and Brückner, 1998). Transcriptomic analysis also revealed that RsaI strongly repressed various key enzymes involved in glucose catabolism pathway such as fructose-biphosphate aldolase (*fba*1), 6-phosphogluconate dehydrogenase 2C decarboxylase (*gnd*), and the thiamine-dependent enzyme transketolase (*tkt*). Additionally, we showed that RsaI repressed the synthesis of fructosamine 3-kinase, which deglycated products of glycation formed from ribose 5-phosphate or erythrose 4-phosphate produced by the pentose phosphate pathway (Gemayel et al., 2007). This enzyme is part of a repair machinery to protect the cells from damages caused by glycation as the results of high glucose concentrations (Deppe et al., 2011). The pentose phosphate pathway is also an alternative route for glucose metabolism, and provides the source of pentose phosphates necessary for nucleotide synthesis. Although it is not known whether RsaI regulated their expression in a direct and indirect manner, base-pairings were predicted between RsaI and the ribosome binding site of *fba*1 and *tkt* mRNAs. The transkelotase is a key enzyme of the pentose phosphate pathway, which requires thiamine diphosphate as a co-factor. Interestingly, the thiamine pathway is also repressed in strain expressing RsaI (Table S4). Although, we have observed very similar pathways that are deregulated by RsaI using the MAPS and RNA-seq approaches, not so many overlaps were identified. It is possible that the conditions of the MAPS approach performed on the wild type strain expressing all the ribonucleases has preferentially enriched the mRNAs that are regulated at the translational level. Therefore, it is tempting to deduct that RsaI would inhibit the synthesis of the major permease of glucose uptake, of enzymes involved in the glycolysis, of unnecessary enzymes involved in detoxification of high glucose concentration, and of the pentose phosphate pathway when glucose concentration decreases.

### RsaI interacts with sRNA containing C-rich motifs

The MAPS approach revealed that several sRNAs (RsaG, RsaD, RsaE) bound significantly to RsaI. We demonstrated here that the second G-track sequence located in the first hairpin domain of RsaI is responsible for the formation of a highly stable complex with RsaG, and a less stable complex with RsaD. Additionally, mRNA targets interacting with the CU-rich domain of RsaI did not disturb the binding of RsaG suggesting that the two functional domains of RsaI are independent. Conversely, preliminary data suggested that the apical loop of the first hairpin of RsaG contain the C-rich motif that is recognized by RsaI, but which is also used to regulate the expression of several mRNAs (Desgranges *et al*., data not shown). RsaG is part of the 3’UTR of *uhp*T mRNA, and its expression was strongly enhanced by the extracellular concentration of G-6P. Under these conditions, the levels of RsaI are much lower than for RsaG (result not shown). These two sRNAs are thus involved in pathways related to the use of the preferred carbon sources. Indeed, RsaI negatively controls glucose uptake when glucose is consumed or absent from the medium while RsaG responded to the extracellular concentration of G-6P. Although the functions of RsaG remained to be addressed, we hypothesized that the sRNA might regulate either the expression of unnecessary genes, or of genes involved in sugar metabolism, or of genes required to protect cells against damages linked to sugar-phosphate uptake and metabolism (Bobrovskyy and Vanderpool, 2014). Noteworthy, RsaI sequence is conserved in the genus of *Staphylococcus* while RsaG is conserved only in the *S. aureus* species.

In addition, the overproduction of RsaI induced numerous changes into the transcriptome of *S. aureus* that resembled the regulon of the two-component system SrrAB, which was demonstrated as the essential system responding to both nitric oxide (NO) and hypoxia (Kinkel et al., 2013). The SrrAB regulon has also been shown to confer to the cells the ability to maintain energy production, to promote repair damages, and NO detoxification (Kinkel et al., 2013). We observed here that RsaI enhanced the expression of genes encoding cytochrome biosynthesis (*qox*ABCD), as well as genes involved in anaerobic metabolism (*pfl*AB, *ldh*1, *foc*A, *adh*), in iron sulfur cluster repair (*scd*A), and most importantly in NO detoxification (*hmp*). These effects might be indirect due to the interaction between RsaI and the SrrAB-dependent sRNAs RsaD and RsaE, which both contained a typical *srr*A site upstream their genes. These two sRNAs present a C-rich sequence that can potentially form base-pairings with the G-track sequences of RsaI (Table 1), but the formation of complexes with RsaI is less efficient than with RsaG. It cannot be excluded that an RNA-binding protein might be required in these cases to facilitate base-pairings. We also do not exclude that the two sRNAs were pooled down because they might share similar mRNA targets with RsaI. RsaD and RsaE are both upregulated by the presence of NO in the cellular medium (Durand et al., 2015; Figure S3D). Although the functions of RsaD requires additional studies, *S. aureus* RsaE was previously shown to coordinate the downregulation of numerous metabolic enzymes involved in the TCA cycle and the folate-dependent one-carbon metabolism (Bohn et al., 2010; Geissmann et al., 2009). Additionally, in *B. subtilis*, the homologue of RsaE called RoxS is under the control of the NADH-sensitive transcription factor Rex, and the Rex binding site is also conserved in *S. aureus rsa*E gene (Durand et al., 2017). Hence, it was proposed that RsaE downregulated several enzymes of the central metabolism under non favorable conditions, and in addition would contribute to readjust the cellular NAD+/NADH balance under stress conditions (Bohn et al., 2010; Durand et al., 2015; 2017). Many of the RsaI-dependent effects, which have been monitored by the transcriptomic analysis, are most probably indirect, and we propose that some of these effects resulted from the interaction of RsaI with sRNAs when the preferred carbon source became scarce.

### Physiological consequences of RsaI regulation

We showed here that several sRNAs in *S. aureus* are part of intricate regulatory networks to interconnect in a dynamic manner various metabolic pathways following sugar metabolism and uptake. The concept that sRNAs are key actors to coordinate the regulation of metabolic enzymes has been largely demonstrated for *Enterobacteriaceae* (reviewed in Görke and Vogel, 2009). These sRNA-dependent regulations often induced significant growth phenotypes in response to the availability of carbon sources and nutrient. For instance, *E. coli* and *Salmonella* SgrS contributed to stress resistance and growth during glucose-phosphate stress (reviewed in Bobrovskyy and Vanderpool, 2014), *Salmonella* GcvB regulon is required for growth if peptides represent the unique carbon source (Miyakoshi et al., 2015), while *E. coli* Spot42 is important for optimal utilization of carbon sources and its overproduction inhibited growth, when succinate was the sole carbon source (Møller et al., 2002). In *S. aureus*, the overproduction of RsaE induced a growth defect, which was partially alleviated by acetate (Bohn et al., 2010). Therefore, the yields of these sRNAs should be tightly controlled during growth in order to adjust the metabolic pathways, to optimize the use of the preferred carbon sources, and to avoid cell damages or metabolites depletion caused by the intracellular production of glucose-phosphate. If RsaI interconnects various metabolic pathways by acting as a sponge sRNA and as a post-transcriptional regulator, what could be its function during the infection process? *S. aureus* has the ability to generate infections through the colonization of many different metabolic host niches. Several studies have shown that both glycolysis and gluconeogenesis systems are mandatory for the infection process, and moreover *S. aureus* appears to be resistant to NO radicals that are heavily produced by the macrophages. Interestingly, glycolysis is an important process that contributes to persist within the macrophages, and to protect the intracellular bacteria against NO (Vitko et al., 2015). However, if the bacteria escape the macrophages or lyse the host cells, *S. aureus* is thought to form aggregates at the centre of highly inflamed and hypoxic abscesses. Under these conditions, the host cells consumed a large amount of glucose to fight the inflammation. Hence, glucose will become scarce for *S. aureus* suggesting that lactate and amino acids derived from the host might be used as the major sources of carbon to enter gluconeogenesis (Richardson et al., 2015). These conditions favored the activation of the two-component system SrrAB, which in turn activates genes required for anaerobic metabolism, cytochrome and heme biosynthesis, and NO radicals detoxification should play an essential role in the survival of the bacteria (Kinkel et al., 2013). Because the CcpA-dependent repression of RsaI is alleviated under conditions where the glucose is strongly reduced, and because many SrrAB-dependent genes are also induced when RsaI is expressed at high levels, it is thus expected that RsaI might also contribute to metabolic adaptations of the cells to the dynamic nature of the host immune environment. Besides, it is also tempting to propose that RsaI might be involved in the dormant state of bacterial cells while environmental conditions are unfavorable (Lennon and Jones, 2011). Interestingly enough, we also observed a RsaI-dependent activation of the expression of the mRNA encoding EsxA, a type VII secreted virulence factor required for the release of the intracellular *S. aureus* during infection (Korea et al., 2014). We also showed here that RsaI regulated the synthesis of the PIA-PNAG exopolysaccharide. Although the precise molecular mechanism is not yet defined, RsaI would modulate the synthesis of the IcaR repressor by binding to its mRNA or by counteracting an activation factor of *ica*R mRNA translation. The synthesis of the PIA-PNAG is tightly controlled according to the metabolic states of the bacterial population. For instance, inactivation of the TCA cycle resulted in a massive derepression of the PIA-PNAG biosynthetic enzymes to produce the exopolysaccharides (Sadykov et al., 2008), and that glucose enhances PIA-PNAG dependent biofilm formation (You et al., 2014), while SrrAB appears as an inducer of PIA-PNAG dependent biofilm (Ulrich et al., 2007).

Although our study shed light on the regulatory activities of this fascinating multifunctional sRNA, there is still much to be learned on how sRNAs can be integrated into the networks regulating the metabolic pathways that are essential for *S. aureus* survival, persistence and invasion within the host.

## MATERIAL AND METHODS

### Plasmids and Strains Constructions

All strains and plasmids constructed in this study are described in Table S1. The oligonucleotides designed for cloning and for mutagenesis are given in Table S2. *Escherichia coli strain* DC10B was used as a host strain for plasmid amplification before electroporation in *S. aureus*. Plasmids were prepared from transformed *E. coli* pellets following the Nucleospin Plasmid kit protocol (Macherey-Nagel). Transformation of both *E. coli* and *S. aureus* strains was performed by electroporation (Bio-Rad Gene Pulser).

The *rsaI* deletion mutant was constructed by homologous recombination using plasmid pMAD in *S. aureus* HG001 and 132 (Arnaud et al., 2004). The deletion comprises nucleotides 2376101 to 2376306 according to HG001 genome (Caldelari et al., 2017). Experimental details are given in supplementary materials.

The vector pCN51::P*rsa*I was constructed by ligating a PCR-amplified fragment (Table S2) containing *rsa*I with 166 pb of its promoter region and digested by *Sph*I and *Pst*I into pCN51 vector digested with the same enzymes. The vector pUC::T7*rsa*I was constructed by ligating a PCR-amplified fragment (Table S2) containing *rsa*I with the T7 promoter sequence and digested by *Eco*RI and *Pst*I into pUC18 vector digested with the same enzymes. Mutagenesis of pUC::T7*rsa*I was performed with Quikchange XL Site-directed mutagenesis (Stratagene) leading to pUC::T7*rsa*I mut1, 2, 3 and 4 (Table S2). To obtain the plasmid pCN51::P3::MS2-*rsa*I, a PCR product containing the MS2 tag fused to the 5’ end of *rsa*I was cloned into pCN51::P3 by digestion of both PCR fragments and of the plasmid by *Pst*I/*BamH*I (Table S2). Plasmids from positive clones were sequenced (GATC Biotech) before being transformed in DC10B, extracted and electroporated into *S. aureus* strains. The plasmids pCN51::P3::MS2-*rsa*I mut2 and mut4 were generated by Quikchange mutagenesis as above. Construction of plasmids pES::rsaI and pES::rsaI mut5 is described in supplementary materials.

### Growth conditions

*E. coli* strains were cultivated in Luria-Bertani (LB) medium (1% peptone, 0.5% yeast extract, 1% NaCl) supplemented with ampicillin (100 μg/mL) when necessary. *S. aureus* strains were grown in Brain-Heart Infusion (BHI), Tryptic Soy Broth (TSB) or Muller Hinton Broth (MHB) media (Sigma-Aldrich) supplemented with erythromycin (10 μg/mL) for plasmid selection. When needed, MHB was complemented with either 0,15% of D-glucose, 0,5% of glucose 6-phosphate, 0,10% of fructose or of xylose (Sigma-Aldrich). NO production was induced by addition of 100 μM Na-diethylamine NONOate (Sigma-Aldrich) as previously described (Durand et al., 2015).

### Preparation of total RNA extracts

Total RNAs were prepared from *S. aureus* cultures taken at different times of growth. After centrifugation, bacterial pellets were resuspended in 1 mL of RNA Pro Solution (MP Biomedicals). Lysis was performed with FastPrep and RNA purification followed strictly the procedure described for the FastRNA Pro Blue Kit (MP Biomedicals). Electrophoresis of either total RNA (10 μg) or MS2-eluted RNA (500 ng) was performed on 1,5% agarose gel containing 25 mM guanidium thiocyanate. After migration, RNAs were vacuum transferred on nitrocellulose membrane. Hybridization with specific digoxygenin (DIG)-labeled probes complementary to RsaI, RsaG, RsaD, 5S, *ccp*A sequences, followed by luminescent detection was carried out as previously described (Tomasini et al., 2017).

### MAPS experiments, transcriptomic and RNA-seq analysis

Crude bacterial extract were prepared and purified by affinity chromatography as previously described (Tomasini et al., 2017). The eluted RNA samples were either used for Northern blot or treated with DNase I prior to RNA-seq analysis. Isolation of tagged sRNAs and the co-purified RNAs was performed in duplicates. The experiments were carried out with the tagged wild type RsaI and two mutant forms (mut2 and mut4). RNAs were treated to deplete abundant rRNAs, and the cDNA libraries were performed using the ScriptSeq complete kit (bacteria) from Illumina. The libraries were sequenced using Illumina MiSeq with a V3 Reagent kit (Illumina), which preserves the information about the orientation of the transcripts and produces reads of 150 nts, which map on the complementary strand. Each RNA-seq was performed at least in duplicates. The reads were then processed to remove adapter sequences and poor quality reads by Trimmomatic (Bolger et al., 2014), converted to the FASTQ format with FASTQ Groomer (Blankenberg et al., 2010), and aligned on the HG001 genome (Caldelari et al., 2017) using BOWTIE2 (Langmead et al., 2009). Finally, the number of reads mapping to each annotated feature has been counted with HTSeq (Anders et al., 2015) using the interception non-empty protocol. To estimate the enrichment values for the MAPS experiment or the differential expression analysis for the transcriptomic experiment, we used DEseq2 (Varet et al., 2016). The statistical analysis process includes data normalization, graphical exploration of raw and normalized data, test for differential expression for each feature between the conditions, raw p-value adjustment, and export of lists of features having a significant differential expression (threshold p-value=0.05; fold change threshold=2) between the conditions. All processing steps were performed using the Galaxy platform (Afgan et al., 2016).

For total RNA extracts and MS2-eluted RNAs, DNase I (0.1 U/μL) treatment was performed 1h at 37°C. The reactions mixtures were then purified by phenol::chloroform:: isoamylalcohol 25:24:1 (v/v) and subsequent ethanol precipitation. RNA pellets were re-suspended in sterile milliQ water. RNA was quantified by Qubit (Life Technologies) and the integrity was assessed with the Bioanalyzer (Agilent Technologies). For transcriptomic, 1 μg of total RNA was ribo-depleted with the bacterial RiboZero kit from Illumina. The TruSeq total RNA stranded kit from Illumina was used for the library preparation. Library quantity was measured by Qubit and its quality was assessed with a Tapestation on a DNA High sensitivity chip (Agilent Technologies). Libraries were pooled at equimolarity and loaded at 7 pM for clustering. The 50 bases oriented single-read sequencing was performed using TruSeq SBS HS v3 chemistry on an Illumina HiSeq 2500 sequencer.

### Preparation of RNAs for *in vitro* experiments

Transcription of RsaI, RsaI mutants and RsaG was performed using linearized pUC18 vectors. PCR fragments containing the 5’UTR of the coding region of selected mRNA targets were directly used as templates for *in vitro* transcription using T7 RNA polymerase. The RNAs were then purified using a 6% or 8% polyacrylamide-8 M urea gel electrophoresis. After elution with 0.5 M ammonium acetate pH 6.5 containing 1 mM EDTA, the RNAs were precipitated in cold absolute ethanol, washed with 85% ethanol and vacuum-dried. The labeling of the 5’ end of dephosphorylated RNAs (RsaI/RsaI mutants) and DNA oligonucleotides were performed with T4 polynucleotide kinase (Fermentas) and [γ^32^P] ATP. Before use, cold or labeled RNAs were renaturated by incubation at 90°C for 1 min in 20 mM Tris-HCl pH 7.5, cooled 1 min on ice, and incubated 10 min at 20°C in ToeP+ buffer (20 mM Tris-HCl pH 7.5, 10 mM MgCl_2_, 60 mM KCl, 1 mM DTT).

### Gel Retardation Assays

Radiolabeled purified RsaI or RsaI mutants (50000 cps/sample, concentration < 1 pM) and cold mRNAs were renaturated separately as described above. For each experiment, increasing concentrations of cold mRNAs were added to the 5’ end labeled wild type or RsaI mutants in a total volume of 10 μl containing the ToeP+ buffer. Complex formation was performed at 37°C during 15 min. After incubation, 10 μL of glycerol blue was added and the samples were loaded on a 6% or 8 % polyacrylamide gel under non-denaturing conditions (300 V, 4°C). Under these conditions where the concentration of the labeled RNA is negligible, the K_D_ dissociation constant can be estimated as the concentration of the cold RNA that showed 50% of binding.

### Toe-printing assays

The preparation of *E. coli* 30S subunits, the formation of a simplified translational initiation complex with mRNA, and the extension inhibition conditions were performed as previously described (Fechter et al., 2009). Increasing concentrations of RsaI were used to monitor their effects on the formation of the initiation complex with *glc*U_2 and *fn3*K mRNAs.

### *In vivo* ß-galactosidase assays

Translation fusions were constructed with plasmid pLUG220, a derivative of pTCV-*lac*, a low-copy-number promoter-less *lac*Z vector, containing the constitutive *rpo*B promoter (Table S1). The whole leader regions of *glc*U_2 (-54/+99), *fn3*K (-33/+99), *tre*B (-23/+99), HG001_01242 (-34/+99), and HG001_02520 (-71/+99) (Table S2) were cloned downstream the P*rpo*B in frame with *lac*Z. The whole gene encoding RsaI with 166 bp of its promoter region was digested with *Pst*I and ligated at the unique *Pst*I of pLUG220::P*rpo*B vector. The final constructs were transformed into the strain HG001Δ*rsa*I. ß-galactosidase activity was measured four times as previously described (Tomasini et al., 2017).

### PIA-PNAG quantification

Cell surface PIA-PNAG exopolysaccharide levels were monitored according to (Cramton et al., 1999). Overnight cultures were diluted 1:50 in TSB-3% NaCl and bacteria were grown at 37 °C. Samples were extracted at 6 h after inoculation. The same number of cells of each strain was resuspended in 50 μl of 0.5 M EDTA (pH 8.0). Then, cells were incubated for 5 min at 100°C and centrifuged 17,000 g for 5 min. Supernatants (40 μl) was incubated with 10 μl of proteinase K (20 mg/ml) (Sigma) for 30 min at 37°C. After the addition of 10 μl of the buffer (20 mM Tris-HCl pH 7.4, 150 mM NaCl, 0.01% bromophenol blue), serial dilutions 1:25 were performed in the same buffer. Then, 10 μl were spotted on a nitrocellulose membrane using a Bio-Dot microfiltration apparatus (Bio-Rad). The membrane was blocked overnight with 5% skimmed milk in phosphate-buffered saline (PBS) with 0.1% Tween 20, and incubated for 2 h with specific anti-PNAG antibodies diluted 1:20,000 (Maira-Litrán et al., 2005). Bound antibodies were detected with peroxidase-conjugated goat anti-rabbit immunoglobulin G antibodies (Jackson ImmunoResearch Laboratories, Inc., Westgrove, PA) diluted 1:10,000, using the SuperSignal West Pico Chemiluminescent Substrate (Thermo Scientific).

## ACKNOWLEDGEMENTS

We thank Thomas Geissmann for helpful advices and discussions, Marie Beaume for the construction of the mutant strain HG001-Δ*rsa*I, and Eve-Julie Bonetti and Anne-Catherine Helfer for excellent technical assistance. We are grateful to Joseph Vilardell for the gift of the plasmid expressing the MS2-MBP, Aurélia Hiron and Tarek Masdek for providing us the HG001-Δ*hptR*S and -Δ*srr*AB mutant strains, and Christiane Wolz for the HG001-Δ*ccp*A and - Δ*cod*Y mutant strain. RNAseq analyses have been partially done using the Roscoff (France) istance of Galaxy (http://galaxy.sb-roscoff.fr/).

